# Glutamate transporter xCT is important for cGAS-dependent interferon responses to DNA and to HSV-1

**DOI:** 10.64898/2026.04.28.721347

**Authors:** Julia Blay-Cadanet, Maria Lange Pedersen, Alice Pedersen, Maja Sønderby Hemberg, Krishna Twayana, Jacob Storgaard, Cecilie S. Bach-Nielsen, Anne L. Thielke, Cecilie Poulsen, Christoph Felix Kollmann, Ensieh Farahani, Michael Lappe, Marie Beck Iversen, Qi Wu, Camilla Bak Nielsen, Mogens Johannsen, Robert Fenton, David Olagnier, Lin Lin, Anne Louise Hansen, Christian Kanstrup Holm

## Abstract

Metabolic reprogramming is a key component of antiviral immunity, yet how metabolite transport regulates innate immune signaling remains incompletely understood. Here, we show that infection with herpes simplex virus 1 (HSV-1) and stimulation with cytosolic DNA induce the cellular export of glutamate via the xCT (SLC7A11) transporter and that inhibition of xCT reduces cellular resistance to viral replication. Mechanistically, xCT inhibition impaired cGAS-STING signaling by reducing DNA-induced cGAMP production, thereby diminishing type I interferon (IFNα/β) responses and downstream induction of interferon-stimulated genes. Interestingly, modulating intracellular glutamate levels through inhibition of other glutamate pathways, e.g., glutaminolysis or glutamate import, also affected cellular IFN responses, suggesting that glutamate is a central control knob for DNA sensing. Finally, we demonstrate that HSV-1 suppresses xCT expression via a mechanism dependent on the immediate early viral protein ICP27, thereby promoting viral replication by limiting cGAS-dependent IFN induction. Together, these findings identify xCT-dependent glutamate transport as a critical metabolic regulator of cGAS-STING-mediated antiviral immunity.

## INTRODUCTION

The interplay between cellular metabolism and antiviral immune responses has emerged as a critical determinant of infection outcome. Viruses frequently reprogram host metabolism to support their replication, while host cells adapt metabolic pathways to restrict viral spread and promote immune activation (*1*). Several classes of viruses induce metabolic rewiring to increase the availability of biomolecules required for viral replication (*2-4*). For example, severe acute respiratory syndrome coronavirus 2 (SARS-CoV-2) disrupts mitochondrial homeostasis and enhances glycolysis to support viral replication and lipid synthesis (*5-7*).

DNA viruses are sensed by cytosolic cyclic GMP-AMP synthase (cGAS). Upon DNA binding, cGAS catalyzes the production of the second messenger cyclic GMP-AMP (cGAMP) (*8-10*). The cytosolic adaptor molecule Stimulator of Interferon Genes (STING) is a sensor of cGAMP, and its binding leads to STING activation and downstream phosphorylation of Tank-binding Kinase 1 (TBK1) and Interferon Regulatory Factor 3 (IRF3) to induce type I interferons (IFNα/β) (*8-12*).

Increasing evidence points to a bidirectional relationship between cellular metabolism and cGAS-STING signaling. Activation of this pathway can reshape metabolic processes such as glycolysis (*13*), while metabolic perturbations can influence DNA sensing by providing immunostimulatory ligands, such as mitochondrial DNA (mtDNA) (*14*), or by affecting cGAS-STING signaling directly (*15*) or indirectly (*16*). Further, post-translational modifications of cGAS by glutamylation have been shown in mice to regulate its enzymatic activity, highlighting a link between metabolic enzymes and DNA sensing (*17*). Despite these advances, the role of amino acid metabolism, and in particular glutamate homeostasis, in regulating cGAS–STING signaling remains poorly understood.

Here, we report that infection of human keratinocytes with the DNA virus herpes simplex virus 1 (HSV-1), as well as stimulation with cytosolic DNA, induces the export of glutamate via the cystine–glutamate antiporter xCT. We demonstrate that xCT-dependent glutamate export is required for efficient cGAMP production, STING activation, and type I IFN responses, thereby promoting antiviral defense. Conversely, inhibition of xCT leads to intracellular glutamate accumulation, impaired cGAS-STING signaling, and increased susceptibility to HSV-1 infection. Furthermore, we find that HSV-1 actively suppresses xCT expression through a mechanism dependent on the immediate early viral protein ICP27, revealing a viral strategy to evade metabolic control of innate immunity. Together, our findings identify glutamate transport as a critical metabolic regulator of DNA sensing and antiviral immunity.

## Results

### Infection with HSV-1 and stimulation with DNA induce xCT-dependent cellular export of glutamate

We first profiled metabolites in the conditioned culture medium of HSV-1-infected human keratinocytes (HaCaT cells) by mass spectrometry. With this approach, we identified multiple metabolites that were significantly altered upon infection, among which glutamate was one of the most prominently increased (**Fig. 1A**). The observation that glutamate metabolism is engaged in response to HSV-1 infection was also noticeable when we re-analyzed a previously published transcriptomic dataset from HSV-1-infected HaCaT cells (*18*). Here, pathway enrichment analysis identified pathways associated with glutamate metabolism among the most affected by the infection (**Fig. 1B-C**).

**Figure 1.**
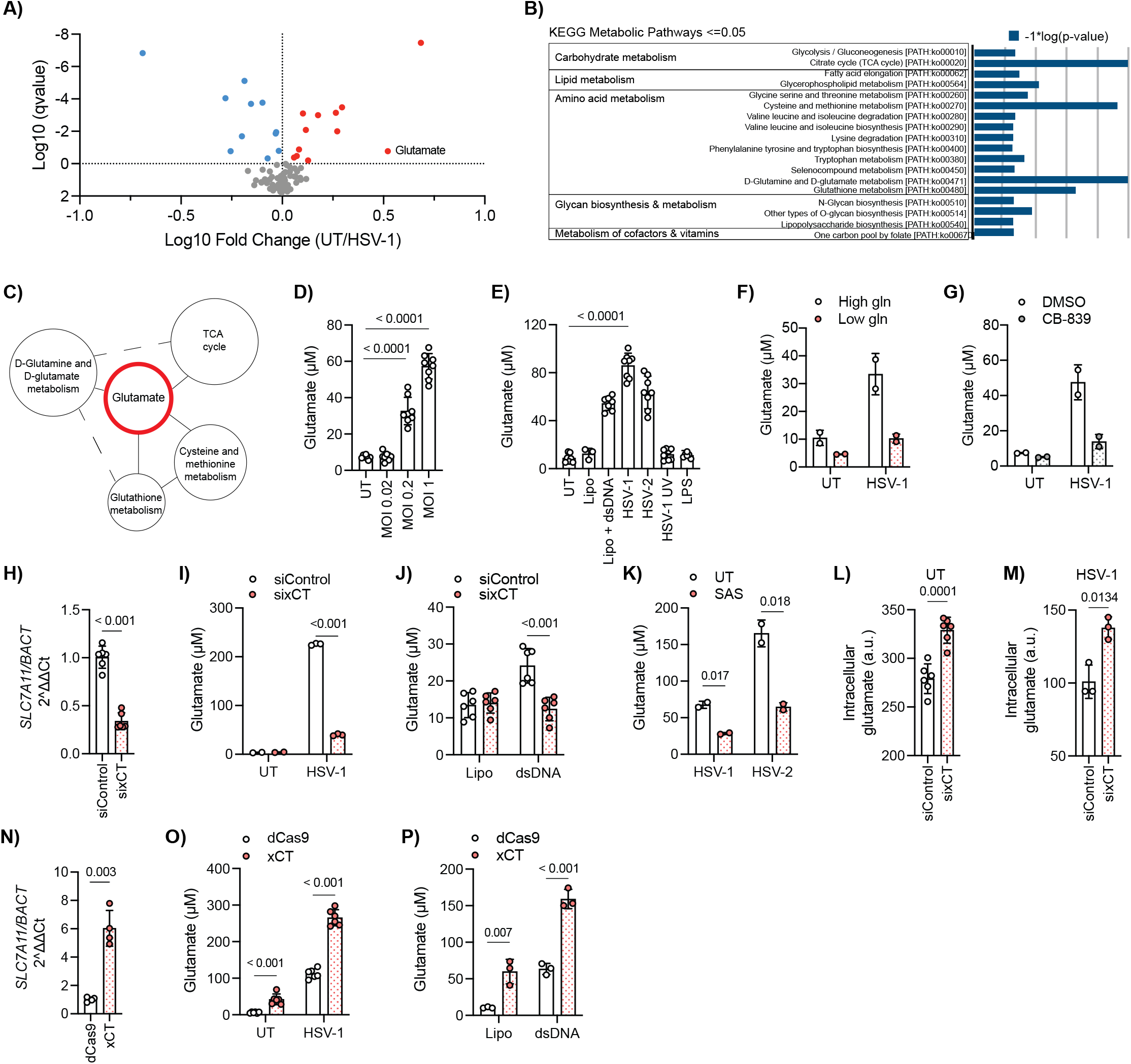
Infection with HSV-1 and stimulation with DNA induce xCT-dependent cellular export of glutamate. (A) Mass spectrometry of conditioned medium from HSV-1–infected HaCaT cells (MOI 0.2, 20 h) identifies glutamate as a major altered metabolite (n = 5). (B–C) KEGG pathway enrichment analysis (Olagnier et al., ref) reveals altered glutamate metabolism in HSV-1–infected cells. (D) Microdialysis measurements show dose-dependent glutamate secretion upon HSV-1 infection (20 h; n = 8, 2 independent experiments). (E) Glutamate release is induced by HSV-1 (MOI 0.2), HSV-2 (MOI 0.2), or cytosolic dsDNA (Lipo + dsDNA) (4µg/mL), but not by UV-inactivated HSV-1 (MOI 0.2) or LPS (1µg/mL), indicating dependence on viral replication (20 h; n = 8, > 3 independent experiments). (F) Glutamate secretion under low-glutamine or standard conditions in uninfected and HSV-1–infected cells (MOI 0.2, 20 h; n = 2, 2 independent experiments). (G) Glutaminolysis inhibition with CB-839 (10 µM) reduces glutamate release in uninfected and HSV-1–infected cells (MOI 0.2, 20h; n = 2). (H) Efficient xCT knockdown by siRNA in HaCaT cells validated by qPCR (72 h; n = 6, > 3 independent experiments). (I–J) xCT knockdown reduces glutamate secretion following HSV-1 infection (MOI 0.2) or cytosolic DNA stimulation (4 µg/mL) (20 h; HSV-1: n = 3, > 3 experiments; dsDNA: n = 6, 2 experiments). (K) Pharmacological inhibition of xCT with sulfasalazine (SAS, 10 µM) decreases glutamate secretion during HSV-1 or HSV-2 infection (MOI 0.2, 20 h; n = 2). (L–M) Intracellular glutamate levels in xCT-depleted cells under uninfected and HSV-1–infected conditions (MOI 0.2, 20 h; n = 6; n = 3, 2 experiments). (N) CRISPR activation induces endogenous xCT expression in HaCaT cells (qPCR; n = 4, 2–3 experiments). (O–P) xCT upregulation enhances glutamate secretion following HSV-1 infection (MOI 0.2) or cytosolic DNA stimulation (4µg/mL) (20 h; n = 6; n = 3, 2 experiments). (a.u. = arbitrary units). Data represent mean ± s.d. of biological replicates. Statistical analyses were performed as described in Methods.

Using microdialysis, we next quantified extracellular glutamate. We observed a dose-dependent increase in glutamate following HSV-1 infection (**Fig. 1D**). A similar response was detected during HSV-2 infection. In contrast, UV-inactivated HSV-1 failed to induce glutamate secretion, indicating that viral replication was required (**Fig. 1E**). Stimulation with cytosolic DNA, but not LPS, also triggered glutamate secretion, suggesting that DNA-sensing pathways contribute to this response. Consistently, kinetic analyses showed a time-dependent increase in both extracellular glutamate and intracellular glutamate levels during HSV-1 infection, indicating enhanced glutamate flux (**Supplementary Fig. 1A-B**).

As intracellular glutamate is primarily generated from glutamine metabolism, we next examined whether glutamine availability influences glutamate secretion. Reducing extracellular glutamine markedly decreased glutamate release under both basal and HSV-1-infected conditions (**Fig. 1F**). Similarly, inhibition of glutaminase (GLS) with CB-839 reduced glutamate secretion during infection, indicating that glutaminolysis is a major source of secreted glutamate (**Fig. 1G**).

The cystine-glutamate antiporter xCT (SLC7A11) is known as the principal transporter mediating glutamate export(*19-21*). We therefore hypothesized that xCT was responsible for glutamate release during HSV-1 infection. Indeed, siRNA-mediated depletion of xCT significantly reduced glutamate secretion following HSV-1 infection and cytosolic DNA stimulation (**Fig. 1H-J**). Pharmacological inhibition of xCT using sulfasalazine (SAS) similarly decreased glutamate release during HSV-1 and HSV-2 infection (**Fig. 1K)**. Conversely, xCT knockdown led to increased intracellular glutamate levels under both basal and infected conditions, consistent with impaired glutamate export (**Fig. 1L-M**).

To further validate this, we employed the CRISPR activation (CRISPRa) system in which nuclease-dead Cas9 (dCas9) recruits RNA polymerase II to the xCT promoter to upregulate endogenous xCT expression (*22*). Increased xCT levels enhanced glutamate secretion under basal conditions and further amplified glutamate release in response to HSV-1 infection and cytosolic DNA stimulation (**Fig. 1N-P**). Together, these results demonstrate that HSV-1 infection and cytosolic DNA sensing increase glutamate metabolism, with increases in intracellular glutamate and secretion, with xCT as the key transporter.

### xCT affects the type I IFN signature

To gain insight into the mechanisms underlying xCT function, we performed bulk RNA sequencing of HaCaT cells following xCT knockdown. This analysis revealed widespread transcriptional changes, with both upregulated and downregulated gene sets (**Fig. 2A**).

**Figure 2.**
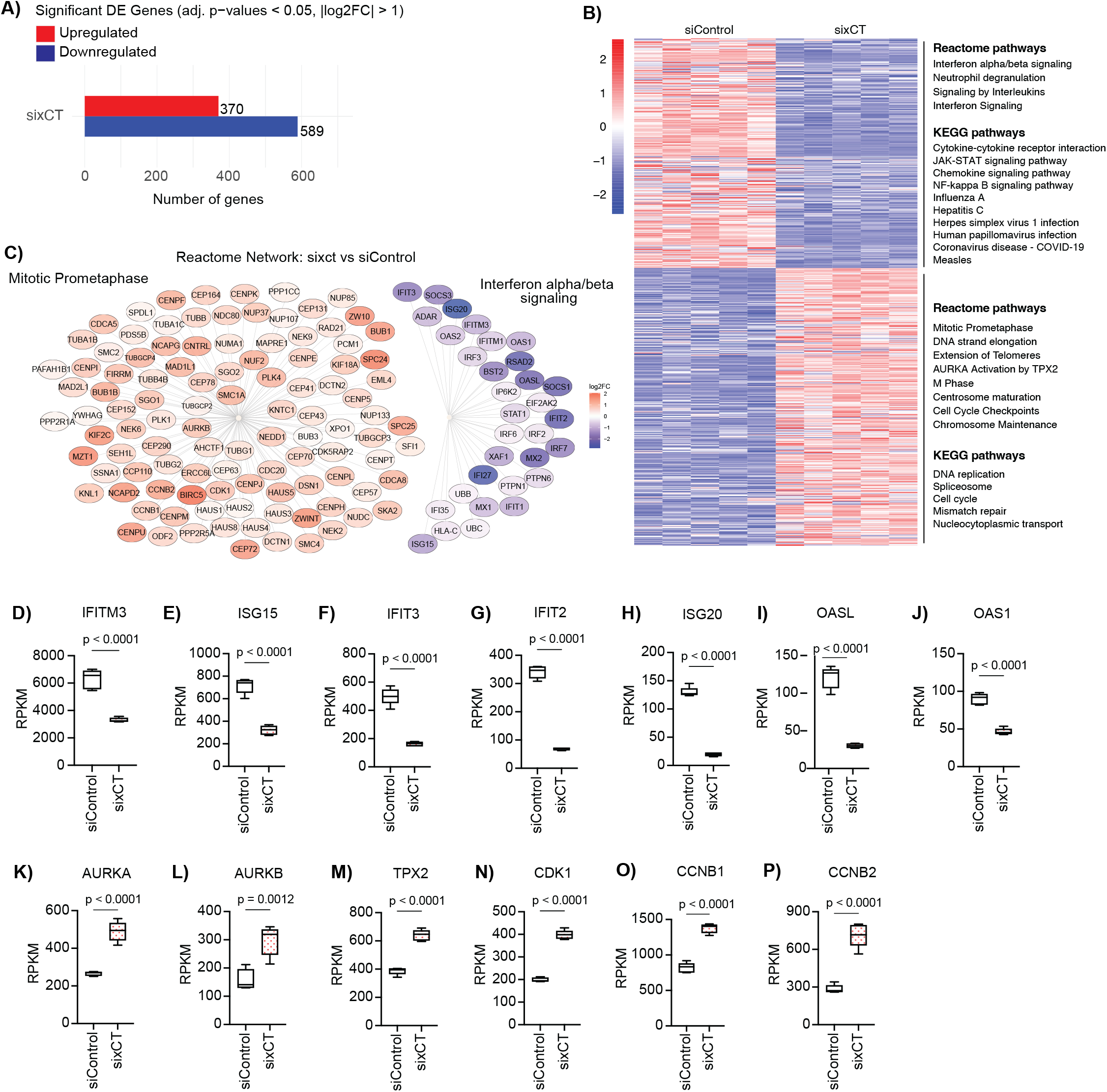
xCT affects the type I IFN signature. RNA-seq analysis of HaCaT cells transfected with control or xCT siRNA for 72 h identifies differentially expressed genes (adjusted p < 0.05, log2FC > 1.0) (n = 5, one experiment). (A) Number of significantly up- and downregulated transcripts. (B) Heatmap of differentially expressed genes with enriched Reactome and KEGG pathways. (C) Word cloud representation of differentially expressed genes. (D-P) RNA-seq read counts for selected genes (RPKM > 50), shown as box plots (mean, minimum, and maximum; n = 5). Data represent mean ± s.d. of biological replicates. Statistical analyses were performed as described in Methods.

Pathway enrichment analysis showed that genes downregulated upon xCT silencing were strongly associated with innate immune and antiviral responses. In particular, interferon alpha/beta signaling (Reactome) and antiviral pathways, including HSV-1 infection (KEGG), were among the most significantly affected (**Fig. 2B**). Notably, xCT silencing led to marked downregulation of multiple interferon-stimulated genes (ISGs), including *IFITM3, ISG15, IFIT2, IFIT3, ISG20, OAS1*, and *OASL* (**Fig. 2C-J**). Conversely, genes upregulated following xCT knockdown were enriched in pathways related to cell-cycle progression, including mitotic prometaphase (Reactome) and DNA replication (KEGG) (**Fig. 2B**). These genes included important regulators of cell cycle progression, including *AURKA, AURKB, TPX2, CDK1, CCNB1*, and *CCNB2* (**Fig 2K-P**). Together, these results indicate that xCT affects the expression of genes associated with anti-viral immunity, the response to type I IFN, and cell cycle progression.

### xCT promotes STING activation and subsequent IFN responses to cytosolic DNA

The transcriptomic analysis indicated that xCT promotes type I IFN responses (**Fig. 2**). To investigate this further, we silenced xCT in HaCaT cells and stimulated cells with cytosolic dsDNA or the STING agonist 2′3′-cGAMP. xCT depletion markedly reduced the induction of *IFNB1* and the IFN-inducible cytokine CXCL10 in response to dsDNA, whereas responses to cGAMP were largely unaffected (**Fig. 3A-C**). Consistently, immunoblot analysis revealed reduced expression of interferon-stimulated genes (ISGs), including STAT1, Viperin, IFIT1, and ISG15, following dsDNA stimulation in cells treated with siRNA targeting xCT (**Fig. 3D**). To assess the contribution of xCT to early DNA-sensing responses, we examined STING activation and downstream IRF3 signaling. Upon dsDNA stimulation, siControl cells displayed robust STING phosphorylation and nuclear accumulation of phosphorylated IRF3 (pIRF3) within 2h. In contrast, sixCT-treated cells showed reduced STING phosphorylation and impaired nuclear translocation of pIRF3, indicating defective activation of the STING-IRF3 axis (**Fig. 3E**). A similar impairment was observed during HSV-1 infection (**Fig. 3F**).

**Figure 3.**
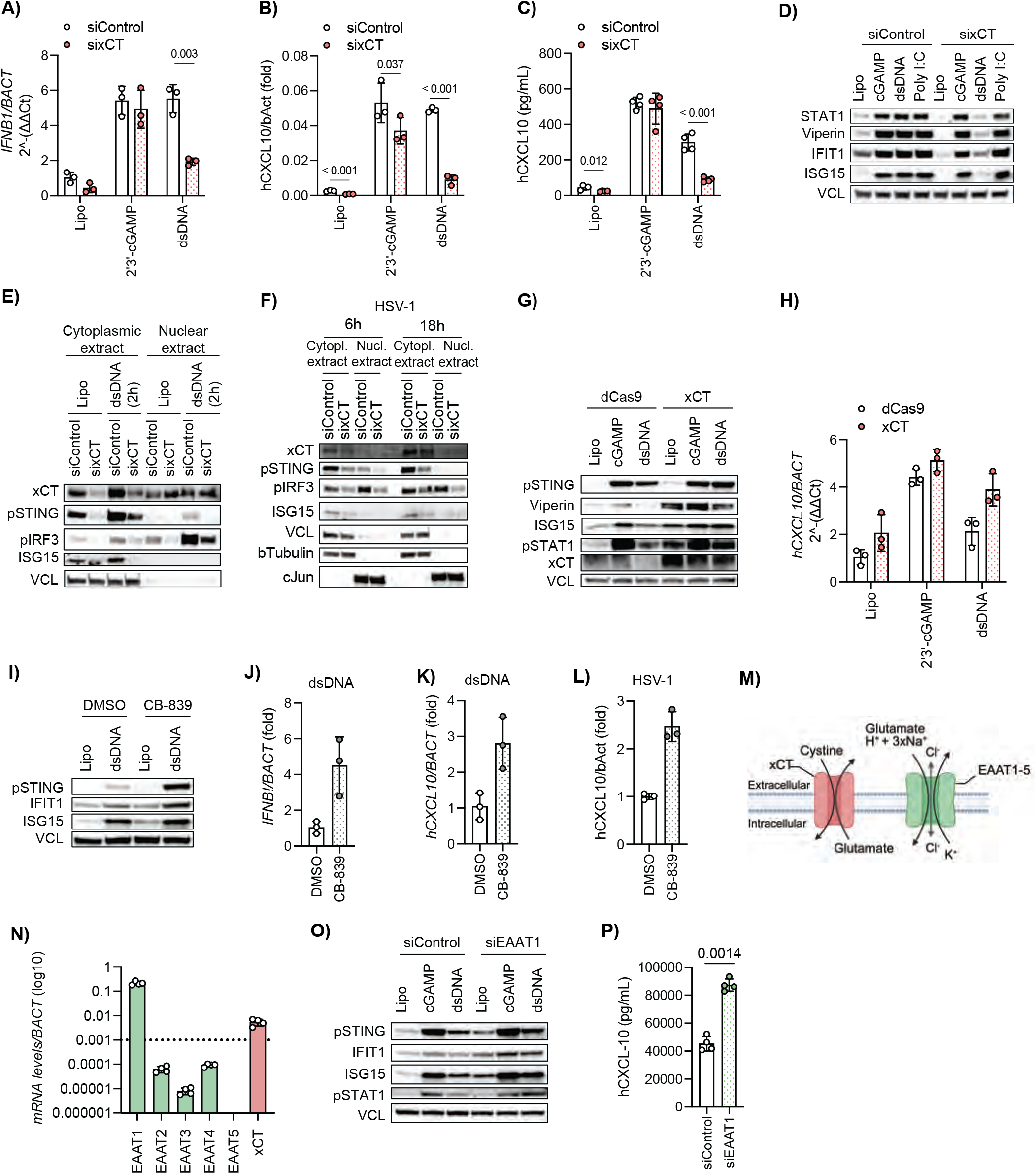
xCT promotes type I IFN response to cytosolic DNA. (A–B) HaCaT cells transfected with control or xCT siRNA for 72 h were stimulated with 2′3′-cGAMP (1 µg/mL), dsDNA (4 µg/mL), or Lipo control for 20 h, followed by qPCR analysis (n = 3, 3 independent experiments). (C) hCXCL10 secretion measured by ELISA in cells treated as in (A–B) (n = 3, 3 independent experiments). (D) STING pathway activation assessed by immunoblotting following stimulation with 2′3′-cGAMP (1 µg/mL) or dsDNA (10 µg/mL) for 20 h. Representative of 3 independent experiments. (E–F) Subcellular fractionation of HaCaT cells transfected with control or xCT siRNA and stimulated with dsDNA (4 µg/mL, 2 h) or infected with HSV-1 (MOI 0.5, 6 h or 18 h), followed by analysis of cytoplasmic and nuclear extracts. (G–H) Interferon responses in CRISPR-mediated xCT upregulation (dCas9–xCT) during stimulation with 2′3′-cGAMP (1 µg/mL) or dsDNA (4 µg/mL) (20 h), assessed by immunoblotting (G) and qPCR (H; n = 3, representative of 2 independent experimetns). (I) Glutaminolysis inhibition with CB-839 (10 µM) alters STING signaling following dsDNA stimulation (4µg/mL, 20 h), assessed by immunoblotting. (J–L) CB-839 treatment (10 µM) reduces type I IFN gene expression following dsDNA stimulation (4µg/mL) or HSV-1 infection (MOI 0.2, 20 h), assessed by qPCR (n = 3). (M) Schematic overview of glutamate transport mechanisms (created with BioRender). (N) Expression of glutamate transporters in HaCaT cells measured by RT-qPCR (n = 4). (O–P) EAAT1 knockdown in HaCaT cells followed by stimulation with 2′3′-cGAMP (1 µg/mL) or dsDNA (4 µg/mL) for 20 h, assessed by immunoblotting (O) and hCXCL10 ELISA (P; n = 4). Data represent mean ± s.d. of biological replicates. Statistical analyses were performed as described in Methods.

We next wanted to test whether forcing increased xCT would improve IFN responses. Forced induction of xCT by CRISPRa enhanced STING phosphorylation and downstream signaling in response to both dsDNA and cGAMP stimulation, as evidenced by increased protein expression of phosphorylated STAT1 (pSTAT1), ISGs Viperin and ISG15, together with increased mRNA levels of *CXCL10* (**Fig. 3G-H**). Notably, xCT induction also increased basal ISG expression, indicating that xCT expression levels also assist in fine-tuning the IFN signature at homeostasis.

Consistent with these findings, modulation of xCT in human monocytic THP-1 cells similarly affected STING pathway activation. Both CRISPR-mediated knockout and CRISPR activation of xCT altered downstream signaling responses to dsDNA and cGAMP stimulation, as assessed by immunoblotting (**Supplementary Fig. 2**), indicating that xCT-dependent regulation of IFN signaling is conserved across cell types.

To determine whether cGAS-STING-dependent IFN responses are broadly affected by metabolic pathways that affect intracellular glutamate availability, we next inhibited glutaminolysis using the GLS inhibitor CB-839. Here, we observed that cells treated with CB-839 displayed enhanced responses to stimulation with DNA, as shown by increased phosphorylation of STING and induction of ISGs such as IFIT1 and ISG15 (**Fig. 3I**). Consistently, CB-839 treatment increased IFNB1 and CXCL10 expression in response to both dsDNA stimulation and HSV-1 infection (**Fig. 3J-L**), indicating that reducing intracellular glutamate promotes DNA-induced IFN-responses.

We next examined whether restricting glutamate uptake would have similar effects. For this purpose, we first assessed the expression levels of known glutamate importers in our keratinocytes. Here, Excitatory Amino Acid Transporter 1 (EAAT1) was the most abundant (**Fig. 3M-N**). Silencing of EAAT1 enhanced STING activation and downstream ISG induction following dsDNA stimulation and increased basal CXCL10 secretion (**Fig. 3O-P**). In contrast, silencing of other glutamate transporters had limited or no impact on pathway activation. Depletion of EAAT2 or EAAT3 by siRNA did not affect IFN responses under stimulated conditions, while silencing of EAAT4 showed only minor or context-dependent effects, potentially because of very low expression levels (**Supplementary Fig. 3**).

Together, these data demonstrate that alterations in cellular glutamate availability, whether through xCT depletion, inhibition of glutaminolysis, or reduced transport via EAAT1, reshape interferon signaling, identifying glutamate metabolism as an upstream regulatory node in DNA-sensing pathways.

### Glutamate transporters xCT and EAAT1 affect cGAMP formation and cGAS protein levels

Building on the observation that xCT regulates STING activation in response to cytosolic DNA but much less so in response to cGAMP (**Fig. 3**), we next examined whether xCT acts at the level of cGAS. We first tested the potential impact of xCT on cGAMP formation. Here, xCT depletion by siRNA markedly reduced cGAMP levels under both basal and dsDNA-stimulated conditions (**Fig. 4A**). In contrast, knockdown of the glutamate importer EAAT1 increased cGAMP production (**Fig. 4B**), indicating that intracellular glutamate availability fine-tunes cGAS activity and is required for efficient DNA-induced cGAMP synthesis.

**Figure 4.**
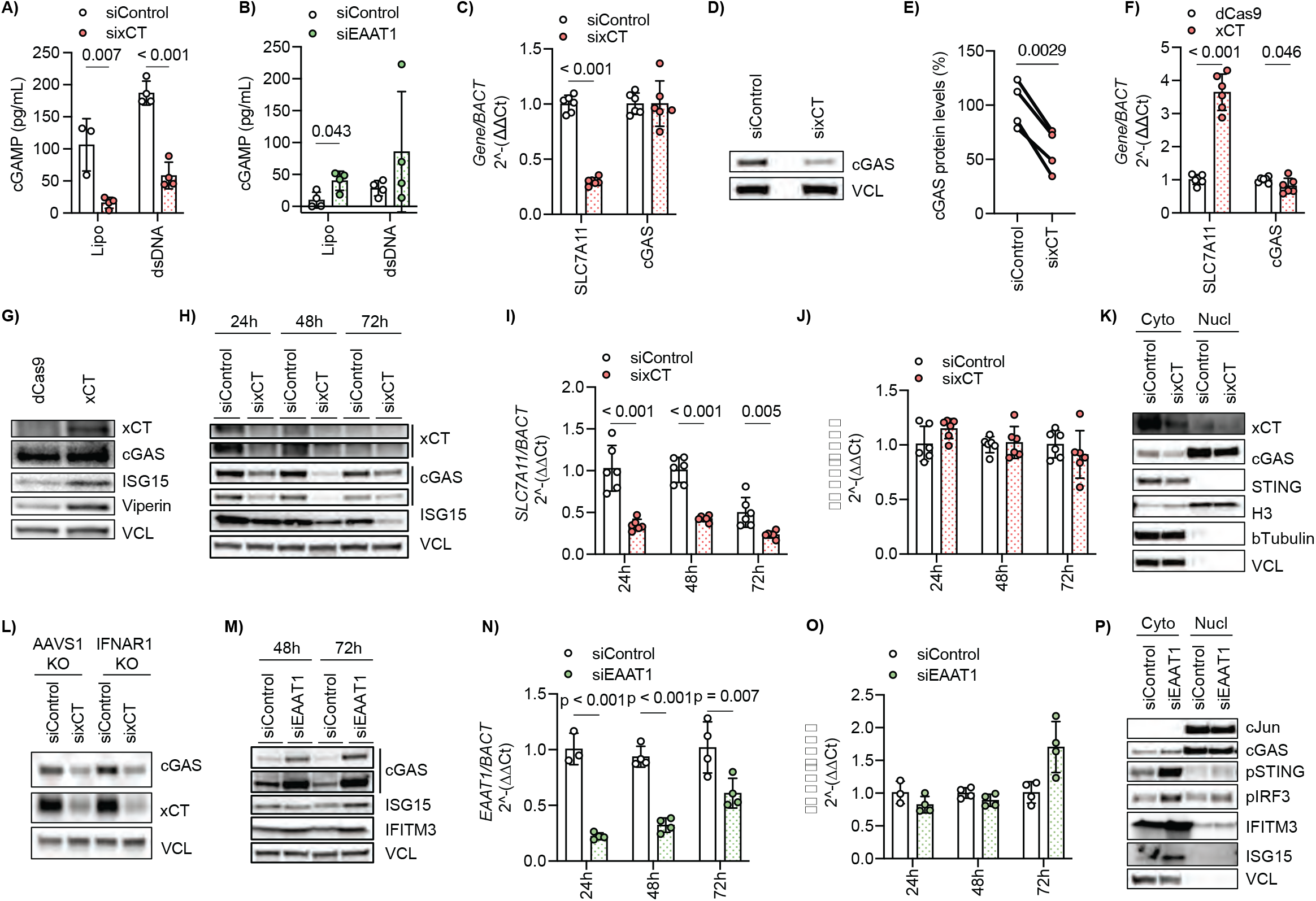
Glutamate transporters xCT and EAAT1 affect cGAMP formation and cGAS expression levels. (A–B) cGAMP production measured by ELISA in HaCaT cells following knockdown of xCT (A) or EAAT1 (B) and stimulation with dsDNA (4 µg/mL, 20 h) (n = 4; representative of 3 (A) or 2 (B) independent experiments). (C) RT-qPCR analysis of cGAS expression following xCT knockdown (72 h) showing no regulation at the mRNA level (n = 6). (D–E) xCT knockdown reduces cGAS protein levels, (E) quantified from immunoblots across 4 independent experiments. (F) RT-qPCR analysis of cGAS expression following CRISPR-mediated xCT upregulation (24 h), showing no regulation at the mRNA level (n = 6). (G) CRISPR-mediated xCT upregulation increases cGAS protein levels. (H) Time-course analysis of cGAS protein levels following xCT knockdown. (I–J) Time-dependent effects of xCT knockdown on gene expression assessed by qPCR (n=6). (K) Subcellular fractionation analysis showing reduced nuclear and cytosolic cGAS following xCT knockdown. (L) Reduced cGAS protein levels are xCT-dependent but independent of IFNAR1 signaling. (M-O) Time-course analysis of cGAS protein (M) and mRNA levels (N-O) following EAAT1 knockdown (n=4). (P) Subcellular fractionation analysis showing increased nuclear and cytosolic cGAS following EAAT1 knockdown. Data represent mean ± s.d. of biological replicates. Statistical analyses were performed as described in Methods.

Given that murine cGAS activity has previously been reported to be regulated by glutamylation, a post-translational modification that can inhibit both DNA binding and enzymatic activity, we next wanted to test whether xCT silencing affected cGAS glutamylation. However, in immunoprecipitation assays, we could not demonstrate any differences in glutamylation upon xCT silencing (**Supplementary Fig. 4A**). Consistently, global glutamylation levels were unchanged as assessed by flow cytometry (**Supplementary Fig. 4B-C**), indicating that xCT does not regulate cGAS through modulation of glutamylation.

We therefore examined whether xCT instead regulates cGAS at other levels. Using siRNA-mediated depletion and CRISPRa-based overexpression of xCT, we measured cGAS expression by RT-qPCR and immunoblotting. While xCT knockdown reduced cGAS protein levels, *cGAS* mRNA remained unchanged, indicating post-transcriptional regulation (**Fig. 4C-E**). Conversely, CRISPR-mediated upregulation of xCT, without affecting transcript levels, had a mild effect on cGAS protein levels, further supporting a role for xCT in controlling cGAS protein abundance (**Fig. 4F-G**). Time-course analysis revealed a progressive decline in cGAS protein following xCT knockdown, whereas mRNA levels remained stable throughout the assay, suggesting that xCT regulates cGAS at a post-transcriptional level (**Fig. 4H-J**). Consistently, subcellular fractionation showed reduced cGAS levels in both cytosolic and nuclear compartments upon xCT depletion, indicating a global reduction in cGAS protein (**Fig. 4K**). Importantly, this effect was independent of type I IFN feedback, as xCT depletion also reduced cGAS protein levels in IFNAR1-deficient cells (**Fig. 4L**), demonstrating that regulation of cGAS by xCT does not require IFN signaling.

We next examined whether modulation of glutamate import via EAAT1 exerts reciprocal effects on cGAS regulation. In contrast to xCT depletion, EAAT1 knockdown increased cGAS protein levels without altering mRNA expression. Time-course analysis revealed a progressive increase in cGAS protein abundance following EAAT1 depletion, whereas transcript levels had no significant changes, consistent with post-transcriptional regulation (**Fig. 4M-P**). Together, these results demonstrate that glutamate transport through xCT and EAAT1 regulates cGAS protein abundance and cGAMP production independently of IFN signaling, thereby controlling activation of the cGAS-STING pathway.

As cGAS function has previously been reported to be regulated during cell division(*23*), we next wanted to investigate the association between suppression of xCT and altered expression of genes associated with the cell cycle, as indicated in the RNAseq analysis (**Fig. 2**). However, despite this transcriptional signature, flow cytometric analysis of cell cycle distribution revealed no detectable differences between control and xCT-silenced cells (**Supplementary Fig. 5**), suggesting that silencing of xCT in this context does not translate into altered cell cycle progression.

### xCT restricts HSV-1 replication

Having established that xCT regulates glutamate availability and DNA-induced IFN-responses, we next examined how xCT affects cellular susceptibility to HSV-1 infection. HaCaT cells were transfected with control or xCT-targeting siRNA for 72 h and subsequently infected with a GFP-expressing HSV-1 strain. Flow cytometry analysis revealed a significantly higher proportion of GFP-positive cells upon xCT silencing, indicating enhanced viral infection using two different virus-cell ratios (**Fig. 5A-B, Supplementary Fig. 6**). Consistently, time-lapse live-cell imaging (Incucyte) confirmed increased viral spread in xCT-depleted cell cultures (**Fig. 5C**). The enhanced susceptibility was further supported by increased viral gene expression, as shown by elevated *vhs* mRNA levels (**Fig. 5D**), as well as higher abundance of viral proteins ICP27 and ICP5 detected by immunofluorescence and immunoblotting (**Fig. 5E-F**). Importantly, plaque assays demonstrated increased production of infectious progeny virus following xCT silencing (**Fig. 5G**). Conversely, CRISPR-mediated upregulation of xCT reduced viral replication, as evidenced by decreased release of infectious virus particles (**Fig. 5H**).

**Figure 5.**
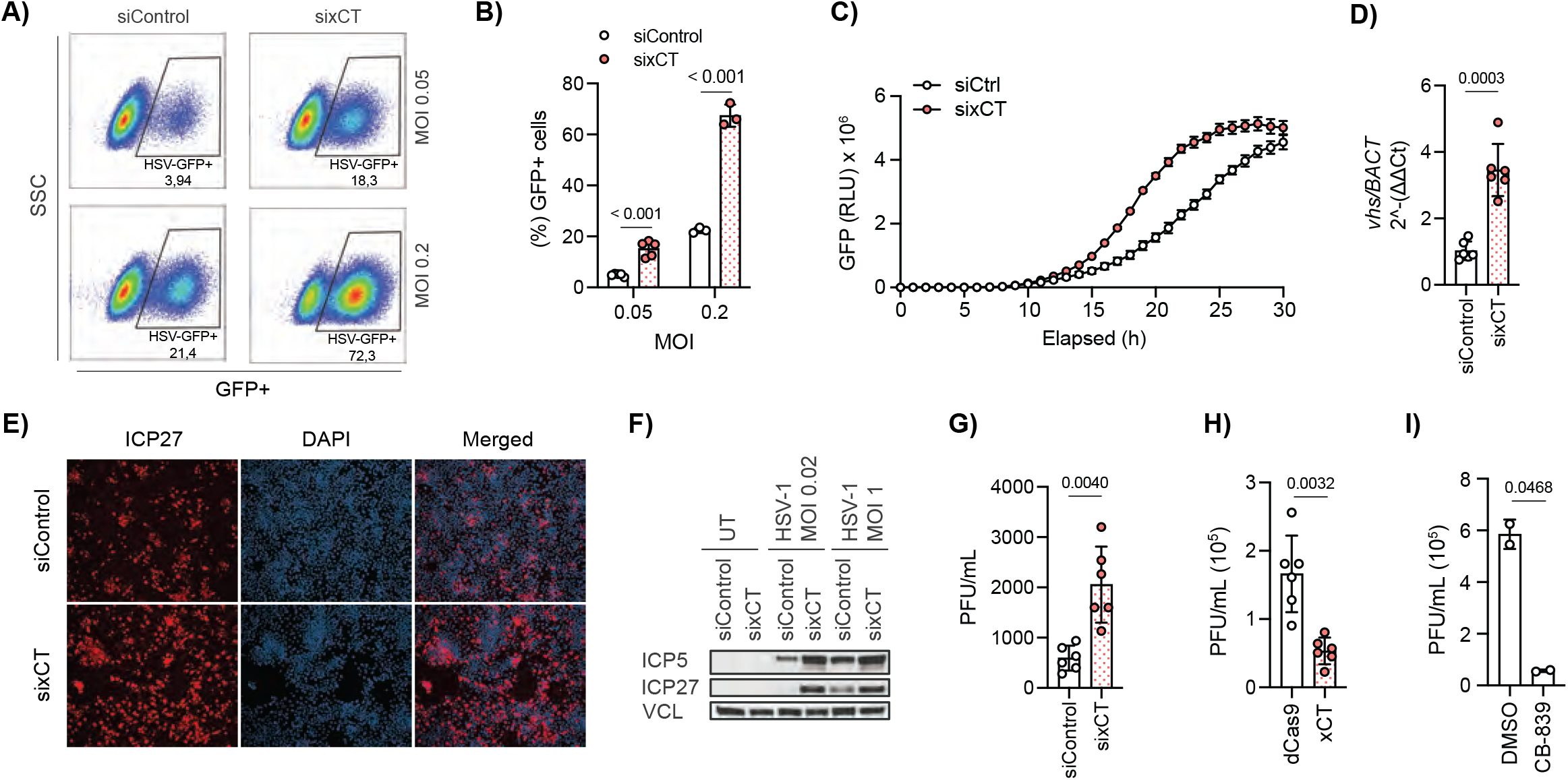
xCT restricts HSV-1 replication by limiting intracellular glutamate accumulation. (A–B) Flow cytometry analysis of GFP-positive HaCaT cells transfected with control or xCT siRNA for 72 h and infected with HSV-1–GFP (MOI 0.05 or 0.2, 20 h). Representative of 2 independent experiments. (C) Time-lapse live-cell imaging (Incucyte) of HSV-1–GFP infection (MOI 0.2) in siRNA-treated cells (72 h) (n = 3; representative of 2 independent experiments). (D) qPCR quantification of viral vhs transcripts following HSV-1 infection (MOI 0.2, 20 h) (n = 6). (E–F) Immunofluorescence and immunoblot analysis of HSV-1 proteins (ICP27 and ICP5) in control and xCT-depleted cells (MOI 0.2, 20 h). (G) Plaque assay quantification of infectious HSV-1 particles released from siRNA-treated cells (MOI 0.2, 20 h). (H–I) HSV-1 replication following xCT upregulation (CRISPR activation) or glutaminolysis inhibition with CB-839, assessed by plaque assay (MOI 0.2, 20 h). Data represent mean ± s.d. of biological replicates. Statistical analyses were performed as described in Methods.

To directly assess the role of glutamate metabolism, we inhibited glutaminolysis using the GLS inhibitor CB-839. Inhibition of glutamate-producing enzyme GLS similarly impaired HSV-1 replication, supporting a functional link between glutamate availability and viral infection (**Fig. 5I**). Together, these results identify xCT as a host factor that promotes type I IFN responses to DNA and infection and restricts HSV-1 replication *in vitro*.

### HSV-1 inhibits xCT expression in an ICP27-dependent manner

HSV-1 encodes multiple mechanisms to evade innate immune responses and promote viral replication. We therefore asked whether HSV-1 suppresses xCT expression to counteract glutamate-dependent promotion of IFN signaling. Infection of HaCaT cells with increasing viral doses revealed a marked, dose-dependent reduction in xCT mRNA levels, indicating that HSV-1 actively represses xCT expression during infection (**Fig. 6A**). As xCT expression is transcriptionally regulated by NRF2, we next examined whether HSV-1 interferes with this pathway. Under basal conditions, chromatin immunoprecipitation revealed NRF2 and RNA polymerase II binding at the xCT promoter, consistent with active transcription (**Supplementary Fig. 7A**). Accordingly, targeting of NRF2 by siRNA reduced xCT expression, and HSV-1-induced glutamate secretion was abrogated in siNRF2-treated cells (**Supplementary Fig. 7B-C**). Further, activation of NRF2 using 4-octyl itaconate (4-OI), Ki696, or siRNA targeting KEAP1, increased xCT expression and glutamate secretion (**Supplementary Fig. 7D-G**). We next examined whether HSV-1 infection alters the capacity of NRF2 agonists to induce xCT expression. Pre-infection of HaCaT cells with HSV-1 abolished the subsequent upregulation of xCT by Ki696 (**Fig. 6B**). Likewise, silencing of KEAP1 by siRNA led to enhanced expression of NRF2 target genes; however, HSV-1 infection markedly reduced the expression of these genes, including HO-1 and xCT (**Fig. 6C**). Overall, HSV-1 appears to antagonize NRF2 signaling and its downstream targets.

**Figure 6.**
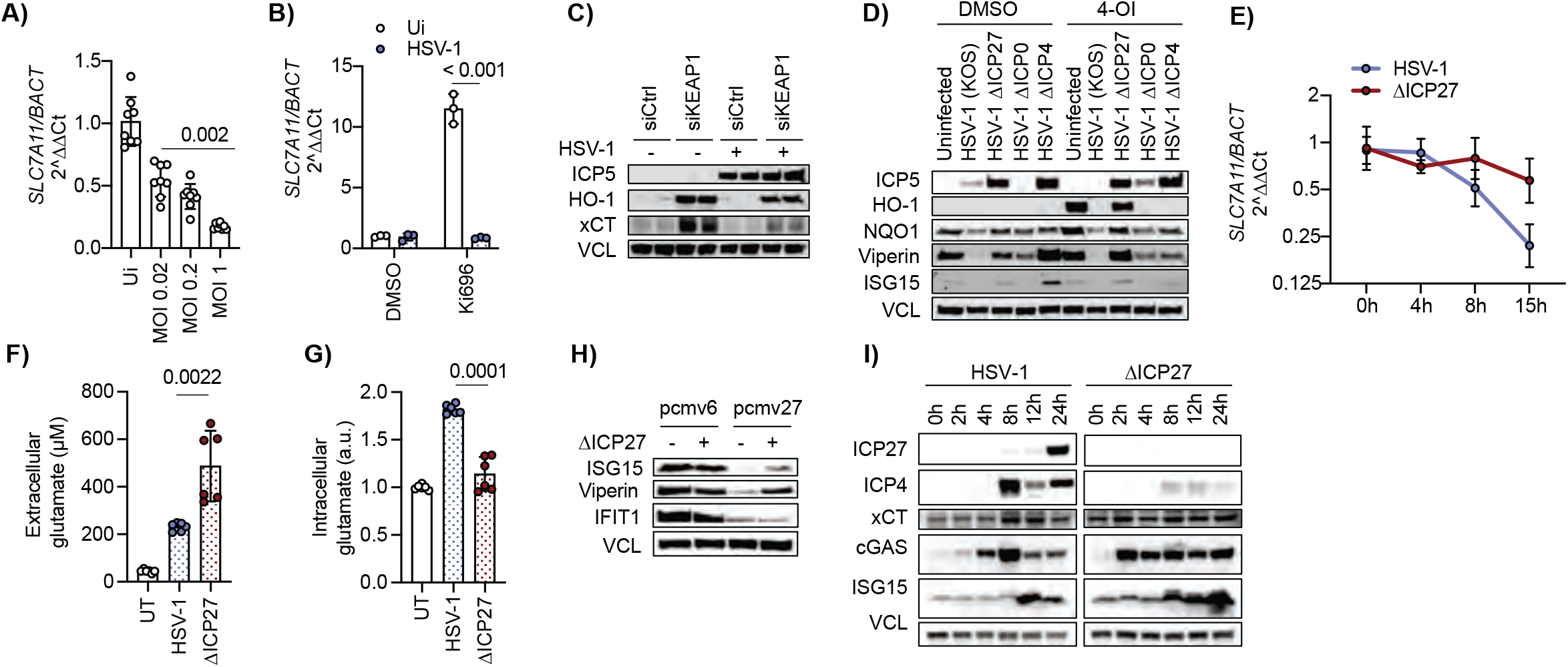
HSV-1 suppresses xCT expression in an ICP27-dependent manner to promote intracellular glutamate accumulation and repress IFN responses. (A) RT-qPCR analysis of xCT expression in HaCaT cells infected with HSV-1 (KOS64) at increasing MOIs for 20 h (n = 8). (B) xCT expression following HSV-1 infection (MOI 0.5, 20 h) and subsequent NRF2 activation with Ki696 (50 µM, 20 h), assessed by qPCR. (C) Induction of xCT by KEAP1 knockdown is suppressed by HSV-1 infection, assessed by immunoblotting. Cells were treated with siKEAP1 or control siRNA for 72 h prior to infection (MOI 0.2, 20 h). (D) Immunoblot analysis of xCT expression following infection with HSV-1 mutants lacking ICP27, ICP0, or ICP4, followed by treatment with or without 4-OI. (E) Time-course analysis of xCT mRNA levels following infection with WT HSV-1 or ΔICP27 mutant (MOI 1), showing loss of repression in the absence of ICP27 (n = 5). (F–G) Extracellular (F) and intracellular (G) glutamate levels in cells infected with WT HSV-1 or ΔICP27 (MOI 1, 20 h) (F: n = 6, 2 independent experiments; G: > 3 independent experiments) (a.u. = arbitrary units). (H) Immunoblotting of ISG15, Viperin and IFIT1 after ectopic expression of ICP27 (20h) modulates following ΔICP27 infection (MOI 2, 20h). (I) Time-course analysis of cGAS protein levels in cells infected with WT HSV-1 or ΔICP27 (MOI 1). Data represent mean ± s.d. of biological replicates. Statistical analyses were performed as described in Methods.

To identify the viral factor responsible for this effect, we analyzed HSV-1 mutants lacking individual immediate proteins. While wild-type HSV-1 suppressed NRF2-dependent gene expression, including HO-1 and NQO1, the ΔICP27 mutant failed to do so, indicating that the immediate early viral protein ICP27 is involved in repressing the xCT-NRF2 axis (**Fig. 6D**). Consistent with this, time-course analysis revealed that wild-type HSV-1 rapidly reduced xCT mRNA levels. In contrast, the ΔICP27 mutant was unable to do so (**Fig. 6E**). Importantly, infection with the ΔICP27 mutant resulted in enhanced glutamate export and reduced intracellular glutamate when compared to infection with wild-type HSV-1 (**Fig. 6F-G**). In line, ectopic expression of ICP27 was sufficient to suppress interferon responses, further supporting a role for ICP27 in regulating this pathway (**Fig. 6H**). Given that xCT silencing affected cGAS protein, we next assessed whether viral targeting of xCT affects cGAS levels. A marked reduction in cGAS protein levels was detected during wild-type HSV-1 infection; however, this effect was not observed with the ΔICP27 mutant (**Fig. 6I**). Together, these findings demonstrate that the immediate early protein ICP27 is involved in suppressing xCT expression, thereby promoting intracellular glutamate accumulation and impairing cGAS-STING-dependent antiviral responses.

## Discussion

Our work identifies a previously unappreciated metabolic response to cytosolic DNA stimulation and HSV-1 infection, in which glutamate is exported via the transporter xCT. Inhibition of xCT in this context increased intracellular glutamate levels and suppressed antiviral responses to DNA. Mechanistically, xCT inhibition reduced the formation of secondary messenger cGAMP by cGAS, which reduced STING/IRF3 activation and subsequently the induction of type I IFNs as well as IFN-stimulated antiviral effectors. These data suggest that intracellular glutamate is an underappreciated regulator of IFN responses to cytosolic DNA. This was supported by the observed increases in IFN-responses when formation of intracellular glutamate was suppressed by inhibition of GLS and when glutamate import was suppressed by inhibition of EAAT1.

Evidence that glutamate can affect cGAS responses is also supported by a previous report, which showed that TTLL4- and TTLL6-mediated glutamylation of murine cGAS regulates its DNA-binding capacity and synthase activity (*17*). Although we confirmed the presence of cGAS glutamylation in our system, we did not observe changes in this modification upon xCT inhibition. Instead, our data indicate that glutamate availability regulates cGAS at the level of protein abundance. Both xCT depletion and EAAT1 knockdown altered cGAS protein levels without affecting mRNA expression, and this regulation was maintained in IFNAR1-deficient cells, indicating that it occurs independently of type I IFN feedback. While the precise mechanisms linking glutamate metabolism to cGAS protein stability remain to be defined, our findings establish metabolic control of glutamate availability as a key determinant of cGAS-dependent IFN responses.

Interestingly, xCT depletion induced a transcriptional signature associated with cell cycle progression, including upregulation of *CDK1* and mitotic regulators. Previous studies have shown that cGAS activity is suppressed during mitosis through CDK1-mediated phosphorylation (*23*), suggesting that cell cycle state may influence DNA sensing. Although we did not detect changes in cell cycle progression, these findings raise the possibility that xCT-dependent metabolic changes intersect with cell cycle-linked regulation of innate immune signaling.

Functionally, inhibition of xCT increased HSV-1 replication and the production of infectious progeny, indicating that this pathway contributes to antiviral defense. In line with this, we found that HSV-1 suppresses glutamate export by downregulating xCT expression through a mechanism dependent on the immediate early viral protein ICP27. These findings expand on the number of ways in which ICP27 targets the STING-signaling pathway and highlight an additional layer by which herpesviruses manipulate host metabolism and antiviral responses (*24, 25*). In this case, rather than directly targeting signaling components, HSV-1 interferes with xCT expression to reshape cellular metabolite homeostasis and dampen IFN responses to DNA.

xCT functions as a cystine-glutamate antiporter, coupling glutamate export to cystine import, which is required for glutathione (GSH) synthesis and protection against oxidative stress. Prolonged inhibition of xCT can induce iron-dependent cell death, also known as ferroptosis, due to impaired GSH production (*26*). However, our observation that the IFN response to DNA was also affected by inhibition of EAAT1 and of GLS, neither of which is associated with ferroptosis, strongly suggests that the effect of xCT inhibition on the IFN response to DNA is not an effect of any engagement of pro-ferroptosis pathways.

Together, our findings demonstrate that human keratinocytes respond to cytosolic DNA and HSV-1 infection by exporting glutamate via xCT, and that this process is required for optimal cGAS-dependent IFN responses and effective control of viral replication. Moreover, HSV-1 targets xCT expression through ICP27 to suppress this pathway, thereby promoting viral replication.

## METHODS

### Cell lines, reagents, and culture conditions

Human immortalized keratinocytes (HaCaT cells) (kindly provided by Søren R. Paludan, Aarhus University, Denmark) were maintained in DMEM (Sigma, #D6429) supplemented with 10% FBS and 1% penicillin-streptomycin-glutamine. THP-1 cells (kindly provided by Bent Deleuran, Aarhus University, Denmark) were maintained in RPMI (Sigma, #R8758) supplemented with 10% FBS and 1% penicillin-streptomycin-glutamine. All cells were maintained in a 5% CO2 incubator at 37 °C. All cell lines were confirmed to be mycoplasma-free. Ki696 was obtained from Sigma (#SML3618-5MG). 4-Octyl-itaconate (4-OI) was synthesized by Thomas B. Poulsen (Aarhus University, Denmark) and dissolved in DMSO. CB-839 was purchased from Cayman (#22038). Sulfasalazine (SAS) was purchased from Merck (#S0083). The viral dsDNA motif 60 mer double-stranded DNA (dsDNA) (tlrl-hsv-60n), the STING ligand 2’3’-cGAMP cyclic (guanosine-(2’-5’)-monophosphate-adenosine-(3’5’)-monophosphate), the dsRNA analog Poly(I:C) (trlr-pic), and the bacterial lipopolysaccharide LPS (tlrl-pb5lps) were purchased from InvivoGen. Intracellular delivery of these reagents was achieved using Lipofectamine 2000 (Thermofisher) diluted in serum-free media (Opti-MEM, Gibco, #31985062) at a 1:1 ratio of the desired concentration.

### Viruses

HSV-1 was propagated and titrated via plaque assay in Vero cells. The viruses used were WT HSV-1 64 GFP+ (strain KOS), HSV-2 (strain 333), HSV-1 ICP27 (KOS), HSV-1 ICP4 (KOS), and HSV-1 ICP0 (KOS). Mutant viruses were kindly provided by Søren R. Paludan (Aarhus University, Denmark). For TCID50/Plaque assay readouts, infected supernatants were removed 1h post-infection, cells were washed, and new media was added to the cells. Cells were then incubated for an additional 17-19h. At 18-20h post-infection, supernatants were collected, and cell pellets were harvested for different analyses. Virus titers in the supernatants were determined by standard plaque assay in Vero cells.

### KEGG pathway analysis

Differentially expressed genes satisfying the threshold of p-value ≤ 0.05 and |(FoldChange)|≥ 1.5 relative to Mock, were used to define host metabolic pathways that are modulated upon HSV-1 infection in HaCaT cells. Gene lists of metabolic pathways were extracted from the KEGG pathway database (Kanehisa et al. 2010), and the overlap between these gene lists and the host differentially expressed genes was assessed by calculating the Z-SCORE relative to 10.000 random trials as a control. The obtained Z-score was used to calculate the p-value. Pathways with a p-value <0.05 were considered significant.

### Short-interfering RNA (siRNA)-mediated knockdown

For short-interfering RNA experiments, HaCat cells were transfected in 6-well plates with 80 pmol of siRNA diluted in OptiMEM (Gibco) and using Lipofectamine RNAiMax (Thermo Fisher) following the manufacturer’s instructions. HaCaT cells were incubated with siRNA for 72 hours before being treated (unless stated differently).

| Target | Assay ID | Target | Assay ID |
| --- | --- | --- | --- |
| NRF2 siRNA (h) | sc-37030 | EAAT1 siRNA (h) | sc-35253 |
| KEAP1 siRNA (h) | sc-43878 | EAAT4 siRNA (h) | sc-41942 |
| xCT/SLC7A11 siRNA (h) | sc-76933 | EAAT3 siRNA (h) | sc-41940 |
| VGLUT1 siRNA (h) | 4392420 | EAAT2 siRNA (h) | sc-35255 |
| Control siRNA (SilencerTM) | 4390844 | VGLUT3 siRNA (h) | sc-42334 |
| Control siRNA-A | sc-37007 | ATG5 siRNA (h) | sc-41445 |

List of siRNA assays used for siRNA silencing.

### Targeted transcriptional activation using the CRISPR-Cas9 system (CRISPRa)

CRISPRa sgRNAs that bind upstream of the transcriptional starting site of the gene xCT were obtained from the Calibrese library (*27*) and purchased from Synthego as chemically modified sgRNAs. The sequences of the sgRNAs are presented in the table below. dSpCas9-VPR mRNA was *in vitro* transcribed from a plasmid that was kindly provided by Rasmus O. Bak (Aarhus University, Denmark) (*28*). Cells were resuspended in Opti-MEM (Gibco, Thermo Fisher Scientific) and electroporated with 95µg/mL mRNA (dSpCas9-VPR) + 50µg/mL of each of the sgRNAs using the 20μL nucleocuvette strip format (Lonza 4D-Nucleofector System (program CM-138-P3)). Electroporated cells (8×10^5 HaCaT cells) were subsequently seeded into a 6-well plate (reaching a total confluency of 80% (1.6×10^6^ cells/well) and cultured overnight before immune stimulation.

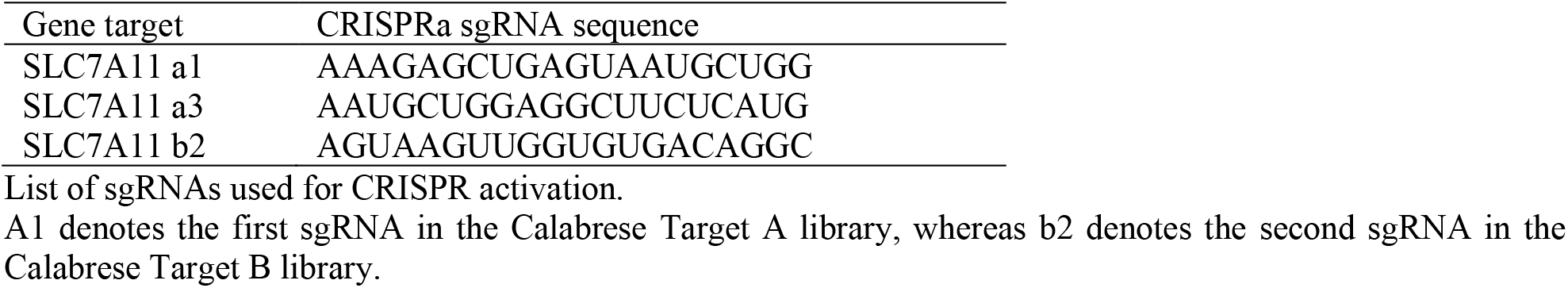

List of sgRNAs used for CRISPR activation.

A1 denotes the first sgRNA in the Calabrese Target A library, whereas b2 denotes the second sgRNA in the Calabrese Target B library.

### CRISPR–Cas9 RNP genome editing (KO)

CRISPR-Cas9-mediated gene disruption was performed in THP-1 and HaCaT cells using ribonucleoprotein (RNP) complexes. Synthetic single-guide RNAs (sgRNAs) targeting *SLC7A11* (*xCT*), IFNAR1, or control locus (AAVS1) were purchased Synthego as chemically modified sgRNA. For RNP assembly, recombinant Cas9 nuclease was incubated with each sgRNA to allow complex formation before delivery. For each gene, genome editing was carried out using two independent sgRNAs (see table below). Preassembled RNPs were mixed with the cells immediately before electroporation and transferred to conductive strip cuvettes compatible with the 4D-Nucleofector platform. Electroporation was performed using the program CM.138-P3, after which cells were returned to culture for recovery.

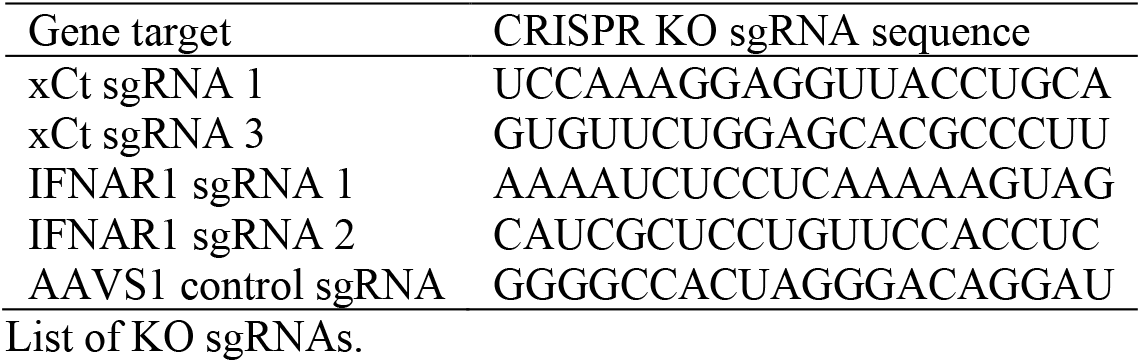

List of KO sgRNAs.

### Glutamate measurements

Cell supernatants were measured on the Micro Dialysis machine ISCUS^flex^ using the Calibrator A (#P000057) and Glutamate Reagent (#P000064). Whole-cell lysates were analyzed with the Glutamate Colorimetric kit from Sigma Aldrich (MAK004-1KT) and the GSH kit from Sigma (CS0260-1KT) following the manufacturer’s instructions.

### Reverse-transcriptase qPCR (RT-PCR)

Total RNA was extracted following the manufacturer’s protocol with the High Pure RNA Isolation Kit (Roche). Gene expression was determined by reverse-transcriptase quantitative qPCR using TaqMan detection systems (Applied Biosystems).

All gene expression results are expressed as arbitrary units relative to the expression of *BACT*.

| Target | TaqMan assay ID | Target | TaqMan assay ID |
| --- | --- | --- | --- |
| ACTB | Hs01060665_g1 | hCXCL10 | Hs00171042_m1 |
| SLC7A11 | Hs00921938_m1 | EAAT1 (SLC1A3) | Hs00904823_g1 |
| HSV-1 | Vi04230116_s1 | EAAT2 (SLC1A2) | Hs01102423_m1 |
| VGLUT1 (SLC17A7) | Hs00220404_m1 | EAAT3 (SLC1A1) | Hs00188172_m1 |
| VGLUT2 (SLC17A6) | Hs00220439_m1 | EAAT4 (SLC1A6) | Hs00192604_m1 |
| VGLUT3 (SLC17A8) | Hs00900423_m1 | EAAT5 (SLC1A7) | Hs00198515_m1 |
| IFNβ1 | Hs01077958_s1 | cGAS | Hs00403553_m1 |

List of TaqMan gene assays used for qPCR.

### ELISA

Cell supernatants were analyzed with hCXCL10 ELISA (R&D Systems).

### Immunofluorescence

Cells were fixed with 4% PFA for 10 min, followed by permeabilization with 0.5% Triton X-100 for 5 min at room temperature. Primary antibody ICP27 (1:100) was added for 1h, followed by further staining with Alexa Fluor TM 647 Chicken anti-mouse IgG (H+L) 2mg/mL #A21463 from Invitrogen (1:500) and DAPI (1 mg/mL conc, #D9542, Sigma-Aldrich). Cells were visualized by the Axio Observer Inverted Widefield, Zeiss - Aarhus University Bioimaging facility.

### Immunoprecipitation assay

Cells were lysed in Pierce IP-lysis buffer (#87788, Thermo) containing 0.5mM NaF and protease inhibitor cocktail, pH 7.4. Lysates were centrifuged at 16,000g for 5 min at 4 °C, and (equalized) supernatants were incubated for 1h at 4 °C with the Anti-FLAG® M2 magnetic beads (#M8823, Merck), followed by immunoprecipitation. Precipitates were isolated with 1%SDS buffer for 10 min at 95 °C and analyzed by immunoblotting.

### Immunoblotting

The cells were harvested, and cell pellets were lysed on ice in ice-cold Pierce RIPA lysis buffer (Thermo Scientific) supplemented with 10mM NaF (a phosphatase inhibitor), 1X complete protease cocktail inhibitor (Roche), and 5 IU/mL benzonase (Sigma). Protein concentration was determined using a BCA protein assay kit (Thermo Scientific), and samples were equalized to the lowest concentration. Whole-cell lysates were denatured for 3 min at 95ºC in the presence of 4X XT Sample Buffer and 20X XT Reducing Agent (Bio-Rad). Samples were loaded and separated by SDS-PAGE on 4-20% Criterion TGX precast gradient gels (Bio-Rad). A marker (Precision Plus protein dual color standard, BioRad) was added to the gel to identify molecular weights. Proteins were transferred from the gel onto PVDF membranes (BioRad) using a Trans-Blot Turbo Transfer system. Membranes were blocked in 5% skim milk (Sigma-Aldrich) at RT in PBS supplemented with 0.05% Tween-20 (PBST) for 1h. Membranes were cut into smaller pieces and incubated overnight at 4ºC with the specific primary antibodies in PBST. After washing in PBST, membranes were incubated with the secondary antibody in PBST 1% milk for 1h at room temperature. After incubation with the secondary antibody, all membranes were washed three times and exposed using either the SuperSignal West Pico PLUS chemiluminescent substrate or the SuperSignal West Femto maximum sensitivity substrate (Thermo Scientific) and detected by an Azure Imaging System 300 (Azure Biosystems).

Secondary antibodies used were Peroxidase-conjugated F(ab)^2^ donkey anti-mouse IgG (H+L) (1:10000) and peroxidase-conjugated F(ab)^2^ donkey anti-rabbit IgG (1:10000) (Jackson Immuno Research).

| Target | Catalog number | Vendor | Dilution |
| --- | --- | --- | --- |
| HSV-1 + HSV-2 ICP5 | ab6508 | Abcam | 1:3200 |
| HSV-1/2 ICP27 | SC-69806 | Santa Cruz Biotechnology | 1:1000 |
| HSV-1/2 ICP4 | ab6514 | Abcam | 1:800 |
| xCT | 12691 | Cell Signaling | 1:1000 |
| xCT | 26864-1-AP | Protein tech | 1:1000 |
| cGAS | 15102 | Cell Signaling | 1:1000 |
| cGAS | 83623 | Cell Signaling | 1:1000 |
| cGAS | 68640-1-Ig | Protein tech | 1:2000 |
| STING | 13647 | Cell Signaling | 1:1000 |
| pSTING | 19781 | Cell Signaling | 1:1000 |
| IRF3 | 11904 | Cell Signaling | 1:1000 |
| pIRF3 | ab76493 | Abcam | 1:1000 |
| TBK1 | 3013 | Cell Signaling | 1:1000 |
| pTBK1 | 5483 | Cell Signaling | 1:1000 |
| STAT1 | 9172 | Cell Signaling | 1:1000 |
| pSTAT1 | 7649 | Cell Signaling | 1:1000 |
| Viperin | 13996 | Cell Signaling | 1:1000 |
| IFIT1 | 14769 | Cell Signaling | 1:1000 |
| ISG15 | 2758 | Cell Signaling | 1:1000 |
| ISG15 | 15981-1-AP | Protein tech | 1:2500 |
| IFITM3 | 59212S | Cell Signaling | 1:1000 |
| IFITM3 | 11714-1-AP | Protein tech | 1:10.000 |
| NQO1 | 3187 | Cell Signaling | 1:1000 |
| HO-1 | 5853 | Cell Signaling | 1:1000 |
| GT335 | AG-20B-0020-C100 | Adipogen | 1:4000 |
| H3 | 4499 | Cell Signaling | 1:1000 |
| c-Jun | 9165 | Cell Signaling | 1:1000 |
| b-Tubulin | ab179509 | Abcam | 1:1000 |
| Vinculin | 18799 | Cell Signaling | 1:1000 |
| Vinculin | V9131 | Sigma-Aldrich | 1:10000 |

Table of antibodies used for immunoblotting.

### Live-cell imaging of GFP reporter virus

Live-cell imaging was performed using the Incucyte® live-cell analysis system (Sartorius). Cells were seeded in appropriate culture plates and transfected with control or xCT-targeting siRNA as described above. Where indicated, cells were treated under the specified conditions and subsequently infected with a GFP-expressing HSV-1 virus. Plates were transferred to the Incucyte system for real-time imaging and maintained under standard cell culture conditions.

Phase-contrast and GFP fluorescence images were acquired at intervals of 1h for up to 72 h using a 10× objective. Cell confluency was determined from phase-contrast images using the Incucyte integrated analysis software. Viral infection was quantified by detection of GFP signal using the green fluorescence channel, and GFP-positive signal was analyzed using the Incucyte analysis software. GFP expression was counted as GFP-positive area or object count per well and, where indicated, normalized to total cell confluency (RCU x µm^2^/image).

### Viral infection and plasmid transfection

Cells were seeded to obtain a confluent cell layer at the time of infection (80-90% confluent) and infected with HSV-1 (MOI=1 or MOI=0.2) at the indicated timepoints. pcmv6 and pcmv27 plasmids were kindly provided by Søren R. Paludan (Aarhus University, Denmark). pcDNA3 N-FLAG cGAS plasmid was kindly provided by Rune Hartmann (Aarhus University, Denmark) and transfected using Lipofectamine 3000.

### Subcellular fractionation

Cellular fractions were isolated following the manufacturer’s instructions for the NE-PER Nuclear and Cytoplasmic Extraction Kit (Thermo Fisher Scientific) (#78833).

### cGAMP quantification

HaCaT cells were treated with dsDNA/(HSV-1) and simultaneously with DMSO or ENPP-1 4e inhibitor (Cayman Chemical #37687). Supernatants and cells were then collected, and cGAMP was measured via a cGAMP ELISA Kit (Cayman Chemical). For cGAMP analysis, whole cell pellets were lysed with M-PER® (Mammalian Protein Extraction Reagent, Thermo Fisher). Cell lysates were analyzed by a 2’3’-cGAMP ELISA kit (Cayman Chemical) according to the manufacturer’s instructions.

### Flow cytometry

The proportion of HSV-1-infected cells was determined by GFP expression at 18-20 h post-infection using flow cytometry (biological duplicates). Cells were harvested and fixed with 1% (vol/vol) formaldehyde (Sigma-Aldrich, F1635) for 15 min at room temperature (RT), washed once with PBS, resuspended in staining buffer (0.5% wt/vol BSA in PBS), and stored cold until analysis. Gating was done on total cells (FSC-H/SSC-H); singlets (FSC-A/FSC-H and SSC-A/SSC-H), and GFP-positive cells (B530 filter/SSC-A) (**Supplementary Fig. 6**). A total of 150,000 cells were acquired per sample. Polyglutamylation levels were analyzed by flow cytometry 72 h after xCT silencing (biological triplicates). Cells were harvested and stained with ZombieNIR Fixable Viability Kit (Biolegend, 423105), washed with PBS, and fixed in 2 % (vol/vol) formaldehyde (Sigma-Aldrich, F1635) for 15 min at RT. Cells were permeabilized with 0.5 % (vol/vol) Triton-X solution for 10 min at RT and washed with staining buffer. Samples were incubated with anti-polyglutamylation modification monoclonal antibody, GT355 (mouse, AdipoGen Life Science, AG-20B-0020-C100, 1:1000) for 30 min, on ice, followed by goat-anti-mouse Alexa Fluor 488 secondary antibody (Invitrogen, A32723, 1:2000) for 30 min on ice. Samples were resuspended in staining buffer and stored cold until analysis. Gating was performed on total cells (FSC-H/SSC-H); singlets (FSC-A/FSC-H and SSC-A/SSC-H); viable cells (R780 filter/SSC-H), and GT355 positive cells (B530 filter/SSC-A) (**Supplementary Fig. 4C**). A total of 100,000 viable cells were collected per sample. Cell-cycle distribution was assessed by flow cytometry 48 h after xCT silencing. Cells were harvested, stained with ZombieNIR Fixable Viability Kit, washed with PBS, and fixed by dropwise addition of ice-cold 70% EtOH while vortexing, followed by incubation on ice for 30 min. Cells were then washed, rehydrated in PBS, and stained with 1 µg/mL 7-aminoactinomycin (7-AAD; AAT Bioquest, 17501) for 30 min at RT. Samples were analyzed immediately using a slow flow rate. Gating and analysis were done on total cells (FSC-H/SSC-H); viable cells (R780 filter/SSC-H); singlets based on the DNA signal (Y667 filter-A/Y667 filter-H), and DNA content, based on 7-ADD intensity on linear scaling (Y667 filter-A) to determine G1, S, and G2 phases using the FlowJo analysis tool fitting to the Dean-Jett-Fox model (**Supplementary Fig. 5B**). A total of 100,000 viable cells were collected per sample. All experiments were analyzed using the NovoCyte Quanteon 4025 analyzer equipped with four lasers (405 nm, 488 nm, 561 nm, and 637 nm) and 25 fluorescence detectors using NovoExpress software v.1.6.2, (Agilent, Santa Clara, CA, USA). Data were analyzed in FlowJo (version 10.8.2 or 10.10.1, Tree Star Inc., Ashland, OR, USA).

### Bulk RNA-seq library preparation

Polyadenylated mRNAs were reverse transcribed using barcoded primers with UMI and template switching to generate full-length cDNA, which was subsequently amplified. For library construction, amplified cDNA was processed using the Illumina-compatible library preparation workflow with a library construction kit (10X Genomics). Final libraries were quality-checked and sequenced on an Illumina NovaSeqX Plus platform.

### Data processing

Raw FASTQ files were aligned to the Homo sapiens reference genome (GRC38, Ensembl 108) using STAR(*29*) (v2.7.11b). Reads were mapped with unique-mapping enabled, and unmapped reads were retained. Gene-level counts were generated using STAR’s GeneCounts/STARsolo module. UMI sequences were extracted and deduplicated. Alignment statistics and count matrices were produced directly by STAR.

### RNA sequencing and data analysis

RNA sequencing data were processed using standardized pipelines at the Bioinformatics Core Facility, Aarhus University. Differential gene expression analysis was performed using DESeq2 (*30*)(v1.46.0). Count data were modeled according to the experimental design, and statistical testing was carried out following the standard DESeq2 workflow. P-values were adjusted for multiple testing using the Benjamini–Hochberg false discovery rate (FDR), and genes were considered differentially expressed at an FDR threshold of <0.05 and an absolute log2 fold change >1.

Pathway enrichment analyses were performed to identify biological processes associated with differentially expressed genes. Over-representation analyses were conducted using curated pathway databases, including KEGG and Reactome categories.

All analyses were performed in R (v4.4.3) using Bioconductor packages on the GenomeDK high-performance computing infrastructure (Aarhus University).

### Mass spectrometry of metabolites

From each sample a 300 µl aliquot of growth media was centrifuged for 5 min at 17.000 g and 200 µl of the supernatant transferred to an Eppendorf tube together with 1.500 µl ice cold MeOH. The solution was vortex mixed for 30 sec and left for 10 min prior to centrifugation for 15 min at 17.000 g. The supernatant was collected and vacuum-centrifuged (SpeedVac, 40 °C) until dry and resuspended in 200 µl Milli-Q water with 0.1% formic acid before LC-MS analysis. A QC sample was prepared by pooling equal volumes of all samples.

Untargeted metabolomics and downstream univariate and multivariate data analysis were done as previously described(*31*). In brief, the LC-MS system consisted of UPLC-QTOF with an ACQUITY I-Class UPLC (Waters Corporation, Milford, MA, USA) equipped with an ACQUITY UPLC HSS T3 column (2.1 mm×100 mm, 1.8 µm, Waters) and coupled to a Bruker maXis Impact QTOF mass spectrometer (Bruker Daltonics, Bremen, Germany). The samples were analysed in random order with an injection volume of 10 µl in ESI− and 2 µl in ESI+. QC samples were injected at the beginning, followed by every fifth sample. The mobile phase consisted of Milli-Q water with 0.1% formic acid (A) and a mixture of MeOH/ACN (1:1 v/v) with 0.1% formic acid (B). The flow rate was set to 0.4 ml min^−1^ at a column temperature of 50 °C with the following 21 min gradient: 0% B (0–2 min), 0–40% B (2–6 min), 40–60% B (6–6.5 min), 60–88% B (6.5–11 min), 88–100% (11–11.5 min), 100% B (11.5–18 min), 100–0% B (18–19.5 min) and 0% B (19.5–21 min). Features were extracted using XCMS (version 4.4.0)(*32*). The centWave algorithm^i^ was used for peak picking with a resolution of 12 ppm and a signal-to-noise threshold of 6. The Obiwarp algorithm was used for retention time correction^ii^. Gap filling was conducted to recover missing signals in the raw data. Features had to be present in at least 80% of the samples within one specific sample group to be considered for further analysis. Isotopes, adducts, and ion source fragments were annotated using CAMERA (version: 1.62.0). Coefficients of variation (CV) of QC samples were calculated to omit features with CV >30%. After analysis, each sample was normalized to the total peak intensity. The areas from the XCMS results were exported to SIMCA (version 18.0.0.372, Sartorius Stedim Data Analytics AB, Göttingen, Germany) for multivariate data analysis. Principal component analysis (PCA) was used as quality control to show the potential cluster trend of the groups and QC samples. Subsequently, orthogonal projections to latent structures discriminant analysis (OPLS-DA) was used to assess the global change between groups. Pareto scaling was applied for both PCA and OPLS-DA. Variable importance in projection (VIP) scores were calculated from OPLS-DA. Fold changes were calculated as mean intensity ratios between groups. Variance equality was tested with an F-test, followed by two-sample t-tests (assuming equal or unequal variances as appropriate). Features were regarded as statistically significant when the P-value <0.05, the fold change >20%, and the VIP score >1. Finally, for identification, m/z value (<5 ppm), retention time, and fragments were compared with an in-house database containing authentic standards.

### Quantification and statistical analysis

All statistical analyses were performed using GraphPad Prism version 10.0. Data are presented as the mean of biological replicates ± standard deviation (s.d.). Statistical significance between groups was determined using an unpaired t-test for normally distributed data, or a two-tailed Mann–Whitney U test for data that did not pass normality testing. Unless otherwise stated, data shown are representative of a single experiment. Most experiments were repeated at least three times with similar results.

## Data availability

The RNA sequencing data are available and deposited to GEO with accession number GSE328494.

## Author contributions

J.B.-C. conceived, designed, and performed experiments, analyzed data, and wrote the manuscript. M.L.P. conducted experiments and analyzed RNAseq data. A.P. conceived experiments. M.S. conceived experiments and analyzed immunofluorescence data. K.T., J.S., A.T.L., and M.B.I. conceived experiments. C.S.B-N. analyzed immunofluorescence images. C.B.N. performed and analyzed mass spectrometry data. Q.W. analyzed mass spectrometry data. C.P. and C.K.F. analyzed Incucyte live-cell imaging data. E.F. and M.L. analyzed RNA-sequencing data. A.L.H. performed flow cytometry experiments, analyzed the data, and contributed to supervision. M.S. and R.F. provided supervision and contributed to data analysis. D.O. conceived and designed experiments, performed data analysis, and supervised the study. L.L contributed to data analysis. C.K.H. designed experiments, analyzed data, supervised the study, and wrote the manuscript. All authors reviewed and approved the final manuscript.

## Acknowledgments

We would like to acknowledge support from Søren R. Paludan for sharing virus strains, and to Rasmus Larsen for his help with processing the RNA-extracted samples for BulkRNAsequencing. We would like to thank the Bioinformatics Core Facility at Aarhus University, especially Jacob Egemose-Høgfeldt.

Flow Cytometry was performed at the FACS Core Facility, Aarhus University, Denmark. Microscopical imaging analysis was performed at the Bioimaging Core Facility, Health, Aarhus University, Denmark.

This work was supported by Novo Nordisk Foundation through a Hallas-Møller Ascending Investigator Grant (NNF21OC0066798), Novo Nordisk project grant in Bioscience and Basic Biomedicine (NNF20OC0064589), Graduate School of the Department of Health Sciences, Aarhus University, Independent Research Fund Denmark (9039-00078B), Hørslev Foundation, Beckett Foundation, Læge Sofus Carl Emil Friis og Hustru Olga Doris Friis’Legat, Leo Nielsen og Hustru Karen Margrethe Nielsens Legat for Lægevidenskabelig Grundforskning, P.A. Messerschmidt & Hustrus Fond, Christian Larsen & Dommer Ellen Larsens Legat and Else og Mogens Wedell Wedellsborgs Fond, the Danish Cancer Society (R279-A16218), Brødrene Hartman Fonden, Einar Willumsens mindelegat, Eva og Henry Frænkels mindefond, Lily Benthine Lunds fonden, Aase of Ejnar Danielsens Fonden, and Frimodt-Heineke Fonden.

---

**Supplementary Figure 1.**
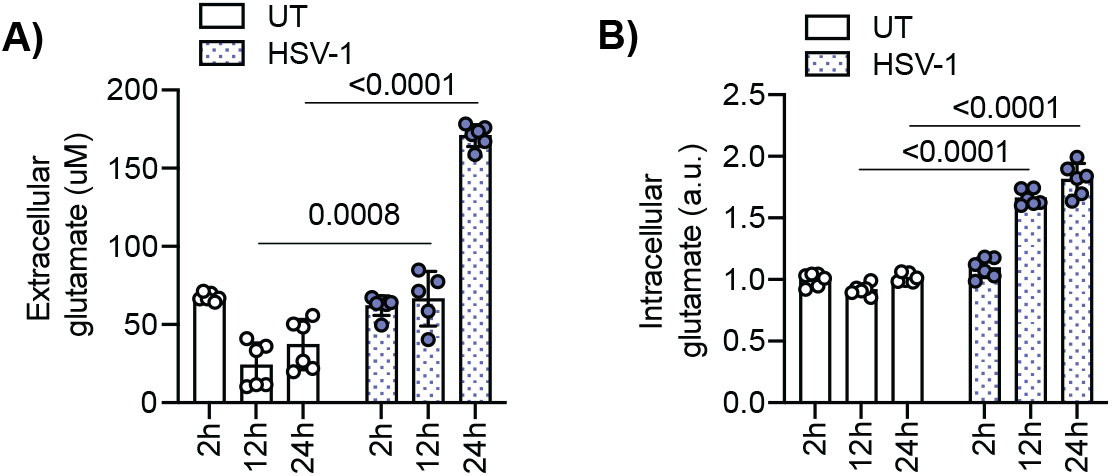
HSV-1 infection and cytosolic DNA stimulation increase extracellular glutamate levels. (A-B) Extracellular and intracellular glutamate measurements in cells infected with HSV-1 (MOI 0.2) over a time course (n = 6). (a.u. = arbitrary units) Data represent mean ± s.d. of biological replicates. Statistical analyses were performed as described in Methods.

**Supplementary Figure 2.**
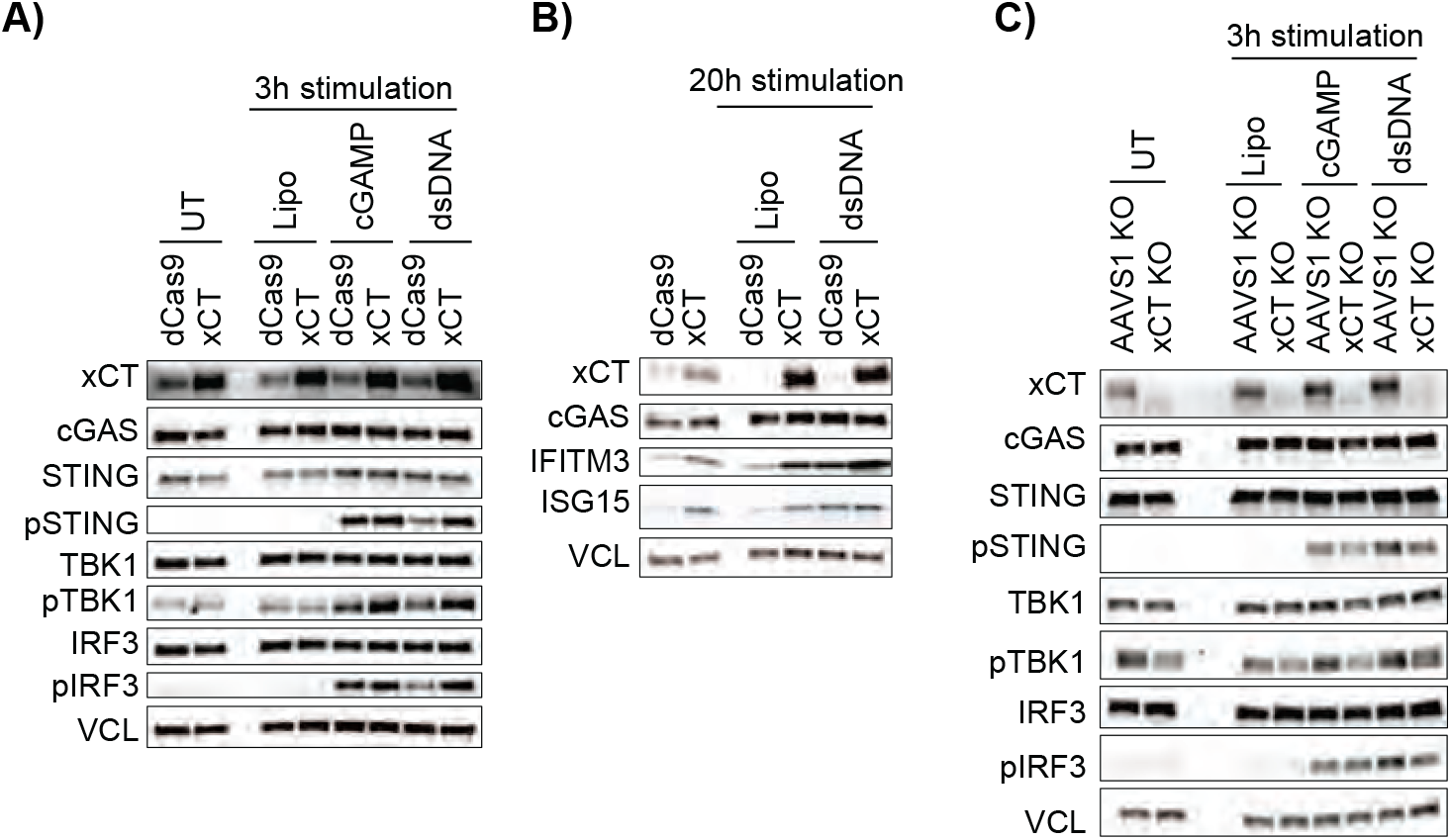
Modulation of xCT alters STING pathway activation in THP-1 cells. (A–B) CRISPR-mediated upregulation of xCT (dCas9–xCT) or control (dCas9) in THP-1 cells followed by stimulation with Lipo, 2′3′-cGAMP (1 µg/mL), or dsDNA (4 µg/mL) for 3 h (A) or 20 h (B), assessed by immunoblotting. (C) CRISPR-mediated knockout of xCT in THP-1 cells followed by stimulation with Lipo, 2′3′-cGAMP (1 µg/mL), or dsDNA (4 µg/mL) for 3 h, assessed by immunoblotting.

**Supplementary Figure 3.**
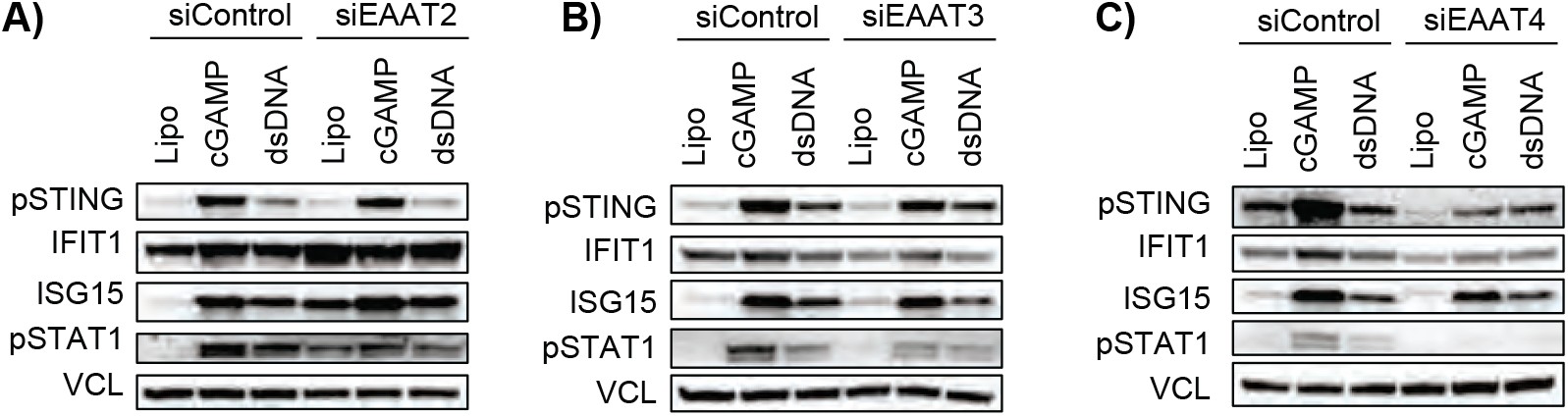
Multiple glutamate transporters modulate cGAS–STING-dependent IFN responses. (A–C) HaCaT cells were transfected with siRNA targeting EAAT2, EAAT3, EAAT4, or control for 72 h, followed by stimulation with 2′3′-cGAMP (1 µg/mL), dsDNA (4 µg/mL), or Lipo control for 20 h. Interferon responses were assessed by immunoblotting.

**Supplementary Figure 4.**
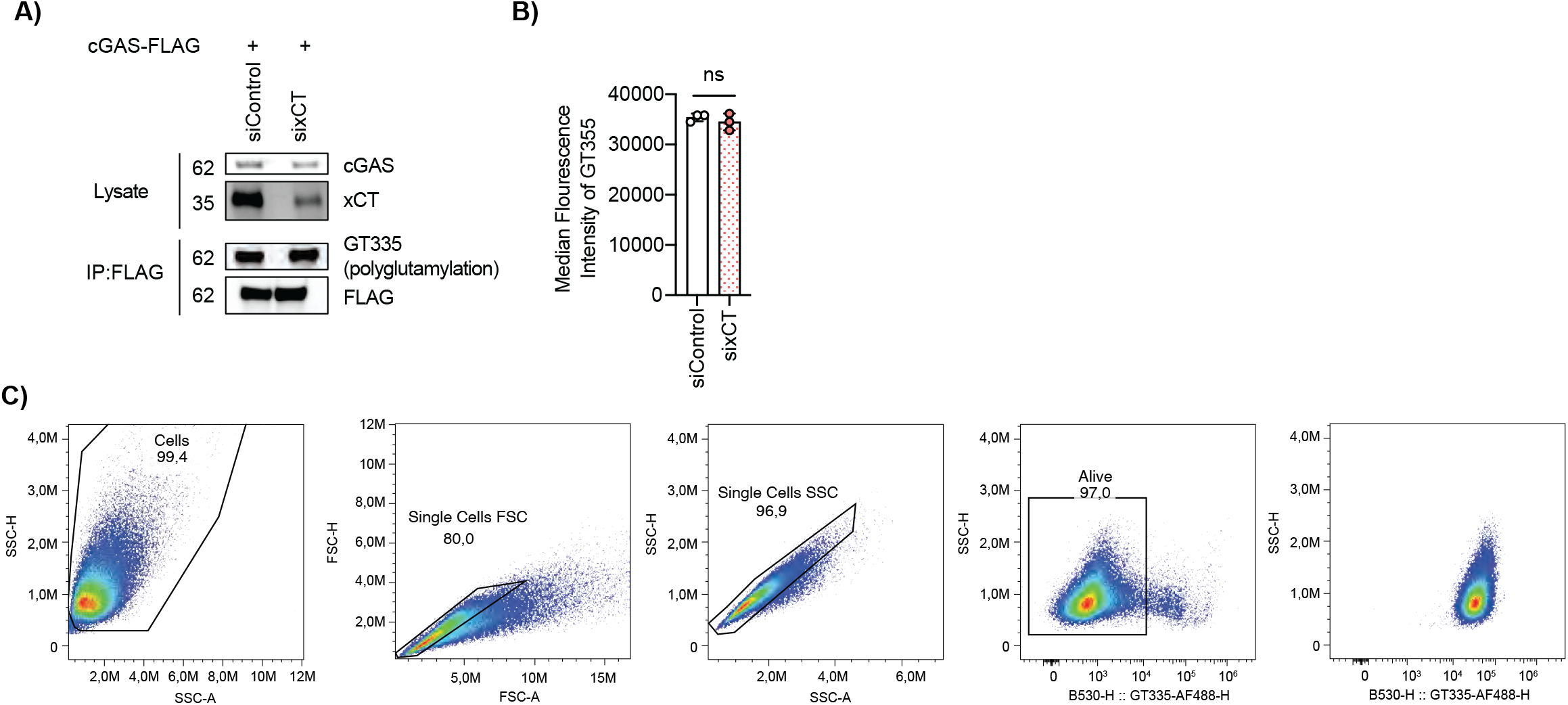
xCT does not regulate cGAS via glutamylation. (A) Immunoprecipitation of cGAS–FLAG in control or xCT-depleted cells shows no difference in GT335 signal. (B) Median fluorescence intensity (MFI) of GT335 measured by flow cytometry is unchanged upon xCT knockdown (n = 3). (C) Gating strategy of B. Data represent mean ± s.d. of biological replicates.

**Supplementary Figure 5.**
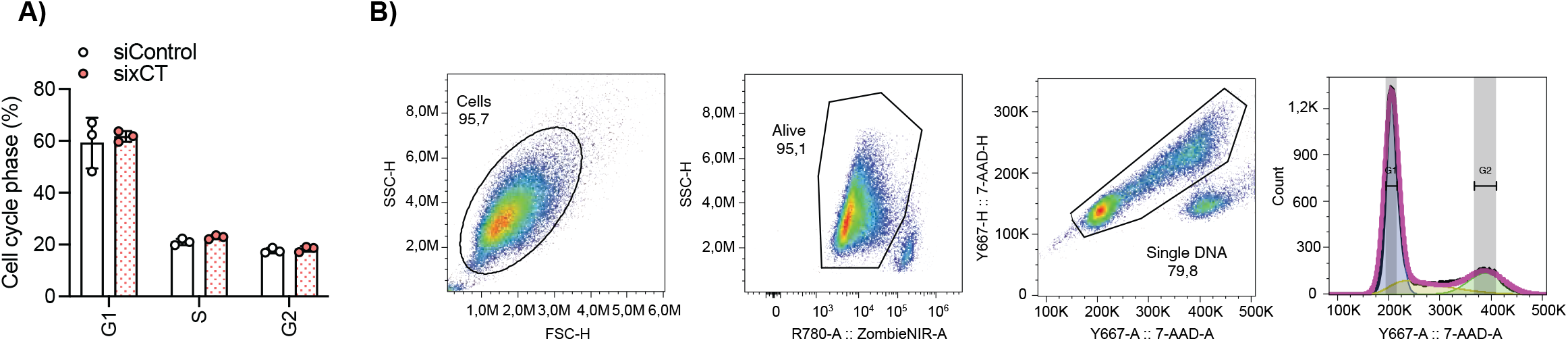
xCT does not affect cell cycle phase. (A) Cell cycle phase is not affeceted after 48h of sixCT vs siControl. (B) Gating strategy of A. Statistical analyses were performed as described in Methods.

**Supplementary Figure 6.**
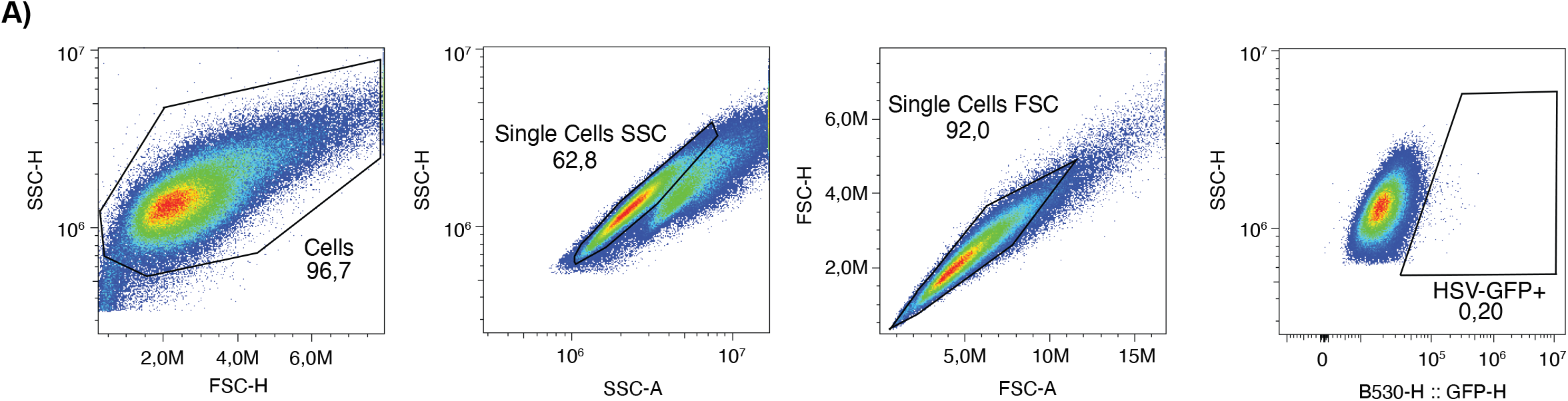
Gating strategy of Fig. 5A.

**Supplementary Figure 7.**
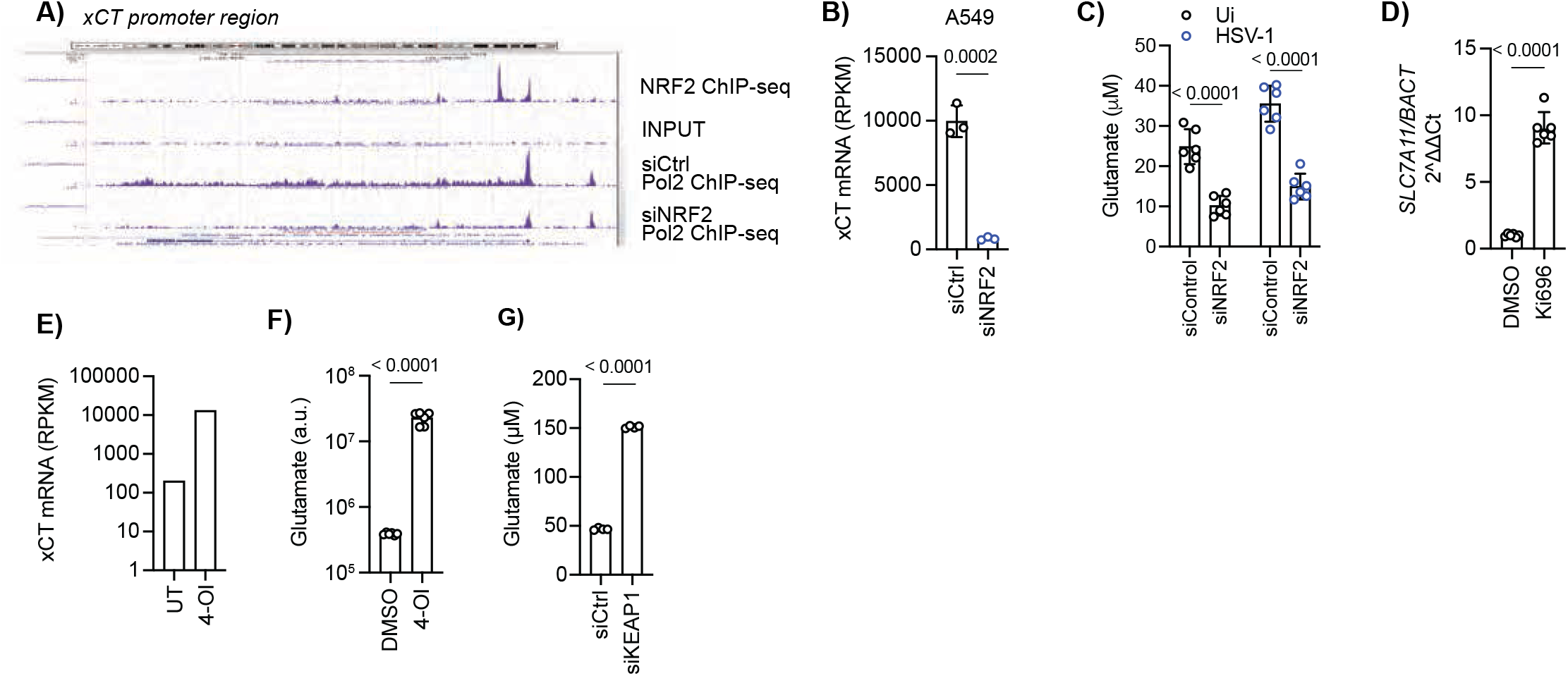
NRF2 controls xCT-dependent glutamate secretion during HSV-1 infection. (A) Chromatin profiling showing NRF2 binding and RNA polymerase II recruitment at the xCT promoter under basal conditions in A549 cells. (B) RNA-seq analysis of xCT expression (RPKM) in A549 cells transfected with control or NRF2 siRNA (n = 3). (C) Reduced glutamate release in NRF2-depleted cells following HSV-1 infection (MOI 0.2, 20 h, n=6). (D) Increased xCT mRNA levels following NRF2 activation with Ki696, assessed by RT-qPCR (n = 6). (E) RNA-seq analysis of xCT expression in HaCaT cells treated with 4-octyl itaconate (4-OI, 125 µM) (showed as mean of n = 3). (F) Glutamate secretion measured by mass spectrometry in cells treated with 4-OI (150 µM, 48 h) (n = 6). (G) Enhanced glutamate secretion following KEAP1 knockdown (72 h; n = 5). Data represent mean ± s.d. of biological replicates. Statistical analyses were performed as described in Methods.

